# Electrical Oscillations of Isolated Brain Microtubules

**DOI:** 10.1101/2020.04.21.054155

**Authors:** Brenda C. Gutierrez, Horacio F. Cantiello, María del Rocío Cantero

## Abstract

Microtubules (MTs) are important cytoskeletal structures engaged in a number of specific cellular activities, including vesicular traffic and motility, cell division, and information transfer within neuronal processes. MTs also are highly charged polyelectrolytes. Recent in vitro electrophysiological studies indicate that different brain MT structures, including two-dimensional (2D) sheets (MT sheets) and bundles, generate highly synchronous electrical oscillations. However, no information has been heretofore available as to whether isolated MTs also engage in electrical oscillations, despite the fact that taxol-stabilized isolated MTs are capable of amplifying electrical signals. Herein we tested the effect of voltage clamping on the electrical properties of isolated non-taxol stabilized brain MTs. Electrical oscillations were observed on application of holding potentials between ±200 mV that responded accordingly with changes in amplitude and polarity. Frequency domain spectral analysis of time records from isolated MTs disclosed a richer oscillatory response as compared to that observed in voltage clamped MT sheets from the same preparation. The data indicate that isolated brain MTs are electrical oscillators that behave as “ionic-based” transistors whose activity may be synchronized in higher MT structures. The ability of MTs to generate, propagate, and amplify electrical signals may have important implications in neuronal computational capabilities.

**Significance Statement:** Microtubules (MTs) are important cytoskeletal structures engaged in a number of specific cellular activities. Recent in vitro electrophysiological studies indicate that different brain MT structures generate highly synchronous electrical oscillations. However, no information is available as to whether isolated MTs also engage in electrical oscillations. In the present study, we provide evidence that non-taxol stabilized isolated MTs generated electrical oscillations with richer frequency spectrum as compared to MT sheets. Thus, structured MT complexes may render more coherent responses at given oscillatory frequencies, suggesting entrainment in combined MT structures. The present study provides to our knowledge the first experimental evidence for electrical oscillations of single brain MTs.

## Introduction

MTs are long hollow cylinders assembled from αβ-tubulin dimer subunits (1–3). It has been long recognized that MTs play an essential role in eukaryote vesicle trafficking and cell division, and in neurons MTs contribute to the development of axons and dendrites (4), and cognitive processing (5) the processing of electrical signals within neurons (6, 7). Previous studies by ours and other groups had determined both experimentally and theoretically the ability of MTs to act as a nonlinear transmission lines capable of electrical signal amplification (8–10). More recent studies also demonstrated that brain MT sheets and bundles generate strong electrical oscillations (10, 11). The electrical behavior of MT sheets was mechanistically consistent with organic electrochemical transistors, where a gate region would drive the synchronous open-close cycles of ion-permeable nanopores that elicit an electrodiffusional circuit, supporting both amplification and self-sustained current-(and voltage-) oscillations (10–12). From a structural viewpoint, this electrical activity may be linked to the nature of the ensemble of the MT subunits. The MT surface forms different structural lattices generated by lateral apposition of protofilaments at least two types of nanopores, at either βαinterdimer interfaces, or βαintradimer interfaces (13). Thus, the electrical phenomenon would implicate changes in nanopore conductance (10). Recent experimental evidence indicated that non-oscillating MT sheets display properties of memristive devices that may underlie the gating mechanism of the MT conductance (12).

In the present study, we explored the ability of single, non-taxol stabilized MTs to handle electrical signals by the loose-clamp configuration of the voltage clamp technique. Isolated MTs generated electrical oscillations at holding potentials different from zero mV. Spontaneous changes in amplitude and frequency were observed in response to both the magnitude and polarity of the driving electrical force, showing qualitatively similar electrodynamic properties as those previously reported for other MT structures (10–13). The data indicate, however, that the spectral oscillatory behavior of isolated MTs may be richer and more chaotic than that of MT sheets, suggesting that MTs organization may induce a strong coherence in the fundamental frequencies in the oscillations. Frequency domain driven electrical information may depend on the structural dynamics of MTs assemblies.

## Materials & Methods

### Preparation of isolated MTs

Commercially available, purified tubulin from bovine brain was utilized in the present study (catalog No. T238, Cytoskeleton, Denver, CO). Tubulin was polymerized with a “hybrid” protocol in the absence of taxol, following loosely general guidelines from the Mitchinson Lab’s online protocols (http://mitchison.med.harvard.edu/protocols/). Briefly, all reactions were conducted in BRB80 solution, containing 80 mM PIPES (1, 4-piperazinediethanesulfonic acid), 1 mM MgCl_2_, 1 mM EGTA, and pH 6.8 with KOH. In some experiments, an aliquot of the tubulin-containing solution was mixed with 2 μl glycerol, 1 mM dithiothreitol (DTT) and 1 mM GTP and incubated for 5 min at room temperature. Isolated MTs were immunochemically labeled with anti α-tubulin antibody (H-300, sc-5546, Santa Cruz Biotechnology Inc) used at 1:500 dilution (11). The secondary antibody was an FITC-tagged bovine anti-rabbit IgG-R (sc-2367, Santa Cruz Biotechnology Inc, CA) used at a 1:1000 dilution. Samples were viewed under DIC and fluorescence microscopy with an inverted Olympus IX71 microscope connected to a digital CCD camera C4742-80-12AG (Hamamatsu Photonics KK, Bridgewater, NJ). Images were collected with the IPLab Spectrum (Scanalytics, Viena, VA) acquisition and analysis software, running on a Dell-NEC personal computer.

### Electrophysiological data acquisition and analysis of isolated MTs and MT sheets

The electronic setup consisted of a conventional patch clamping amplifier (Axopatch 200B, Molecular Devices, Sunnyvale CA), directly connected to the MT via a saline-containing patch pipette, as previously reported (10, 11). Patch pipettes were made from soda lime capillary tubes (Biocap, Buenos Aires, Argentina) with 1.25 mm internal diameter and a tip diameter of ~4 μm. Pipette tips most often rendered resistances in the order of 4-15 MΩ in an “intracellular” KCl solution containing, in mM: KCl 140, NaCl 5, EGTA 1.0, and Hepes 10, adjusted to pH 7.18 with KOH. To initiate the experiment, an isolated MT (Fig. 1b) was identified by both DIC and fluorescence, approached by the patch pipette, and attached by light suction from the pipette tip (Fig. 1a, c). The voltage clamp protocol included steady steps at different holding potentials (gap-free protocol), from zero mV. Voltage corrections under “loose clamp” configuration were conducted as described in (11). Electrical signals were acquired and filtered at 10 kHz, digitized with an analog-digital converter (Digidata 1440A, Molecular Devices) and stored in a personal computer with the software suite pCLAMP 10.0 (Molecular Devices), also used for data analysis. Sigmaplot Version 11.0 (Jandel Scientific, Corte Madera, CA) was used for statistical analysis and graphics.

**Fig. 1:**
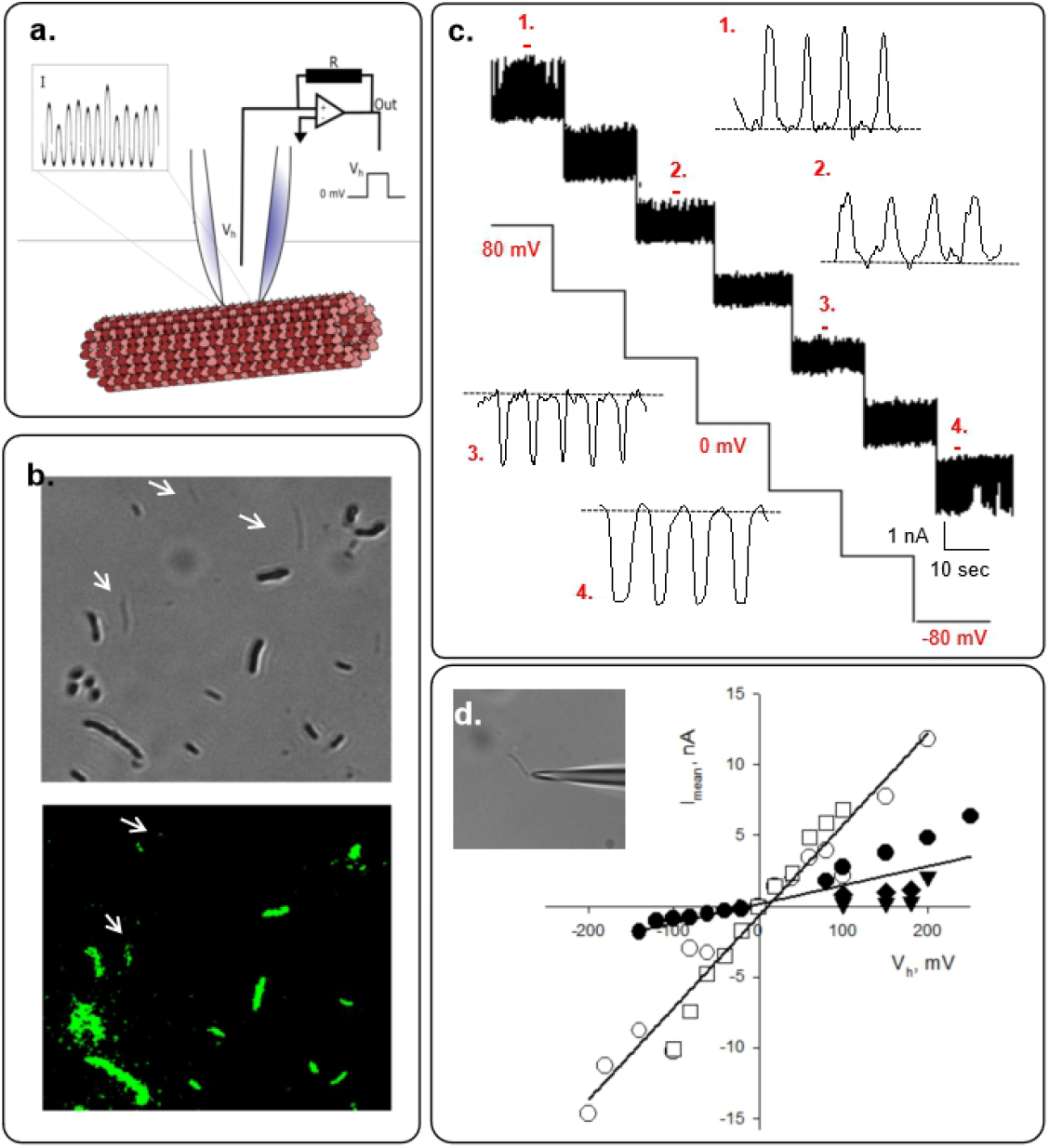
Experimental setup and electrical recordings from isolated brain microtubules. (**a**) Schematics of the patch-clamp configuration used in the study. (**b**) DIC (*Top Panel*) and fluorescent images (*Bottom Panel*) of MTs prepared as indicated in Methods (x60). White arrows shows isolated MTs (**c**) Electrical recordings at different holding potentials (±80 mV) as indicated. Expanded tracings show electrical oscillations of regions numbered “1” through “4”. Applied voltages represented the driving potentials at the patch clamp amplifier. (d) Mean current-to-voltage relationship obtained in symmetrical KCl. Open and closed symbols represent individual experiments for high and low conductance, respectively (n = 5). Inset shows a pipette approaching an isolated MT.

### Other current analyses

Electrical tracings shown throughout the study were unfiltered data. Average currents were obtained by integration of one-second tracings expressed as mean ± SEM values, where (*n*) represented the number of experiments analyzed for a given condition. Power spectra of unfiltered data were obtained by the Fourier transform subroutine of Clampfit 10.0. Poincaré diagrams were constructed by the time delay (*τ*) approach, where the lag time *τ* was chosen arbitrarily at 2*f*, and *f* was the data acquisition sampling frequency.

### Solutions and chemicals used in electrophysiological experiments

All reagents were obtained from Sigma-Aldrich (St Louis MI, USA).

## Results & Discussion

The present study was conducted to obtain electrical recordings from isolated MTs polymerized with a protocol that avoided the use of the MT stabilizer taxol, which inhibits electrical oscillations in other MT preparations (10), and may explain previous failure to observe electrical oscillations in isolated MTs (8, 9). The “loose” patch clamp configuration was used under symmetrical ionic conditions, namely identical saline composition in both patch pipette and bathing solution (see Materials & Methods). Isolated MTs were visualized and identified under combined DIC and fluorescent microscopy by exposure to both a primary anti α-tubulin antibody and an FITC-fluorescent secondary antibody (Fig. 1b, c).

Experiments were conducted with small tipped (4 μm^2^) pipettes. Apposition of the pipette tip onto an isolated MT (Fig. 1a, b) only increased the seal resistance in 1.24 ± 0.29 MΩ reaching to 12.84 ± 5.20 MΩ, n = 14, Median 6.50 MΩ, and a range of 2.83 M Ω to 63.3 MΩ. The approach rendered a type of loose patch configuration as previously used with brain MT bundles (Cantero 2018), and in contrast to the high seal resistances usually obtained with MT sheets (18). Isolated MTs initially voltage-clamped at zero mV showed no electrical activity in any successful experiments (n = 14). MTs displayed spontaneous, self-sustained electrical oscillations (20/29) (Figs. 1a & 2) in direct response to the magnitude and polarity of the electrical stimulus. However, spontaneous oscillations showed changes in regime, particularly by changes in polarity of the holding potential (Fig. 1c). Mean oscillatory currents were linear with respect to holding potential (Fig. 1d), showing at least two different conductances of 13.4 ± 2.5 nS and 64.8 ± 3.2 nS (n = 5). After loose-patch correction, the average change in conductance was 160.8 ± 7.6 nS (n = 3), which is much higher than previously reported for more complex MT structures (18, 19). Fundamental frequencies were often seen at 13, 29, 38, 48, and 90 Hz (Fig. 2a & b). At a given holding potential, oscillations spontaneously changed with time, both in amplitude and the pattern of frequencies (Figs. 2a & b). An increase in the magnitude of the holding potential increased the amplitude of the oscillations, evidencing more frequencies and more complex oscillatory behavior (Fig. 3).

**Fig 2:**
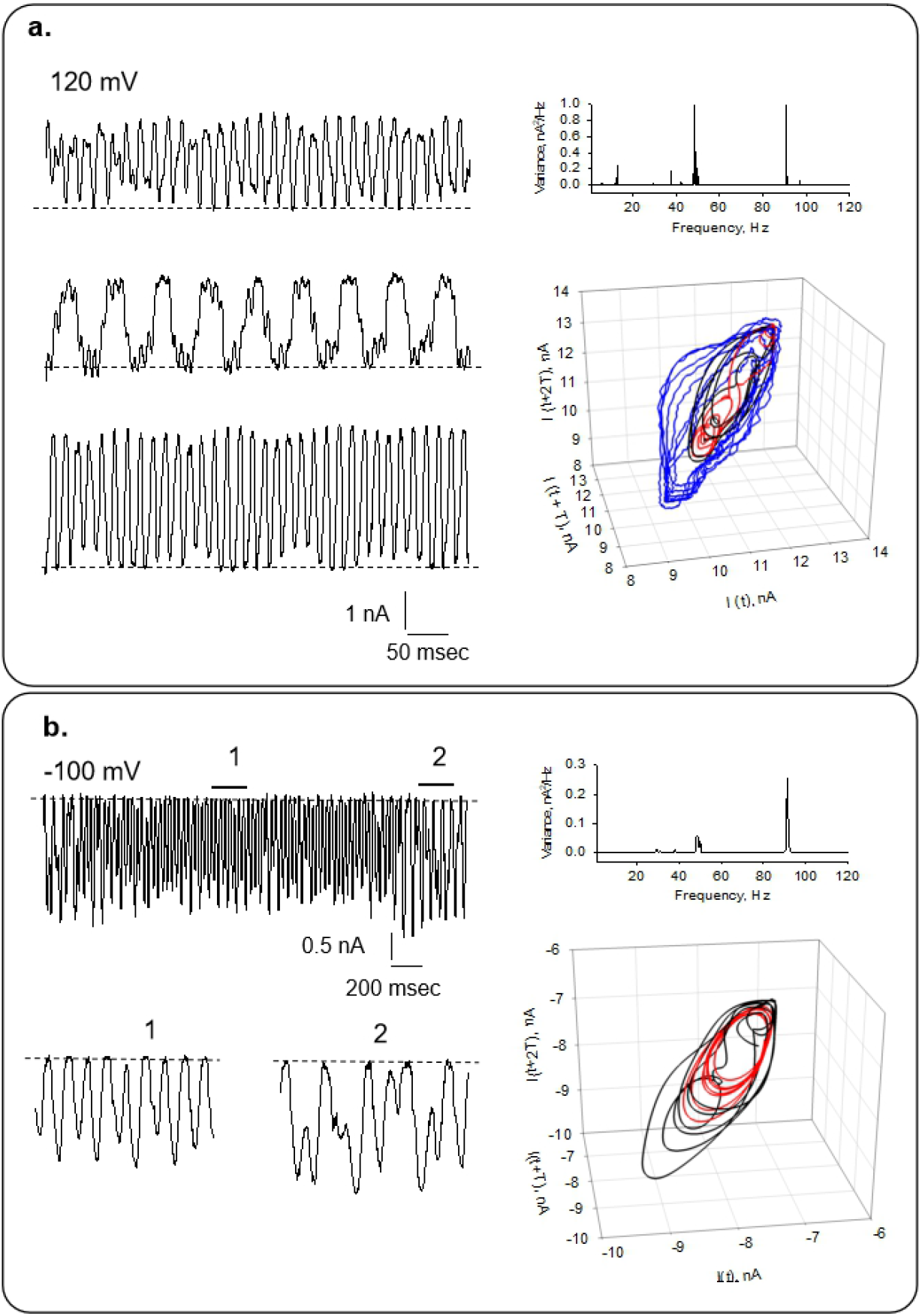
Current oscillations of voltage-clamped brain microtubules at holding potentials of different polarity. (**a**) *Left.* Example of changes observed in oscillatory behavior from an isolated MT at 120 mV. Tracings represent different instances of the same record. *Right*. (*Top*) The panel shows the power spectrum from oscillatory currents shown in tracings on Left. (*Bottom*) 3D Poincaré map of time derivatives from the tracing shown in (**a**). Black, red and blue lines correspond to Top, Middle and Bottom tracings, respectively. (**b**) *Left.* Representative changes in the pattern of oscillations at bottom panels (1 & 2) at −100 mV. *Right*. (*Top*) Power spectrum from oscillatory currents corresponding to the tracing shown on the Left. 3D Poincaré map of time derivatives of tracings (*Bottom*). Red and black lines correspond to “1” and “2”, as indicated in the electrical recording.

**Fig 3:**
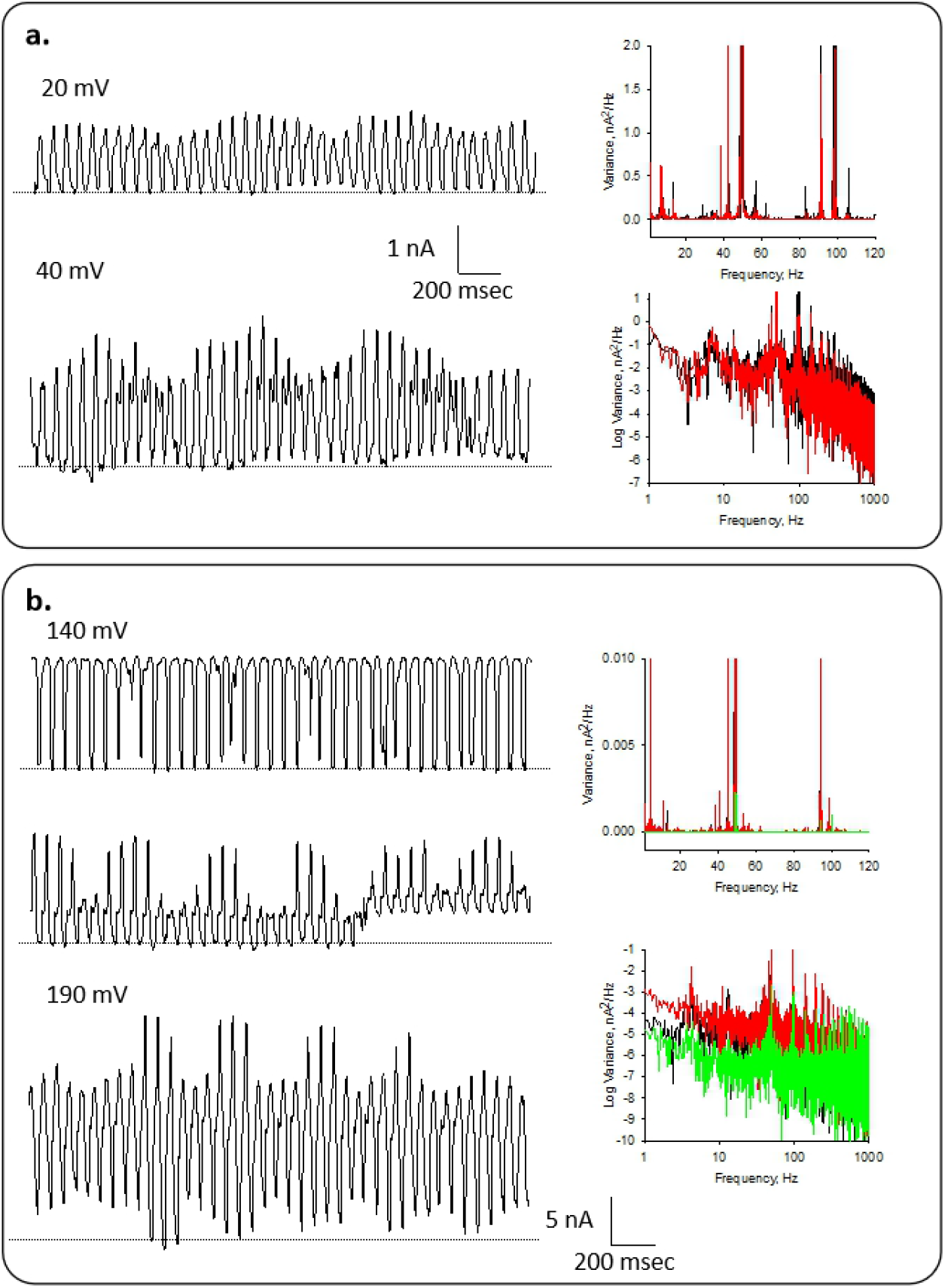
Different patterns of current oscillations at different holding potentials. (**a**) *Left*. Representative recording of an isolated MT at 20 mV and 40 mV, as indicated. *Right.* Linear-linear (Top) and Log-Log (Bottom) plots of Fourier power spectra obtained from tracings on *Left*. Black and red lines correspond to 20 mV and 40 mV, respectively. (**b**) Representative recording of voltage-clamped isolated MT at 140 mV and 190 mV, as indicated. At 140 mV, tracings show different instances of the same record. *Right.* Linear-linear (Top) and Log-Log (Bottom) plots of Fourier power spectra obtained from unfiltered current responses of tracings on *Left*. Black, green and red lines correspond to 140 mV (top tracing), 140 mV (middle tracing) and 190 mV, respectively.

To further explore the rich oscillatory behavior of isolated MTs, similar experiments were conducted under identical conditions, including the same pipette before and after attachment to either an MT sheet, or isolated MT present in the same preparation (Fig. 4). Clearly, the unattached pipette showed a largely white noise spectrum with contaminant line frequency peaks at 50 and 100 Hz (Fig. 4a & d), while MT-attached spectra displayed colored peaks of relevant frequencies in the oscillatory behavior. Clear differences were observed in the MT sheets (n = 4), where it was found fundamental frequencies at 38 and 90 Hz as confirmed by Fourier transformation (Fig. 4b & d). In all cases, the frequency peaks were richer in isolated MTs, as compared to their corresponding MT sheet (Fig. 4c & d). Thus, MTs in the larger surface area of the MT sheet (with tighter seal as well), displayed more coherent behavior in terms of fundamental frequency peaks observed. Thus, an interesting difference between isolated MTs and MT structures became evident, namely, a richer oscillatory behavior of isolated MTs that drastically changed by assembling into flat surfaces showing fewer but stronger oscillatory peaks (Fig. 4).

**Fig 4:**
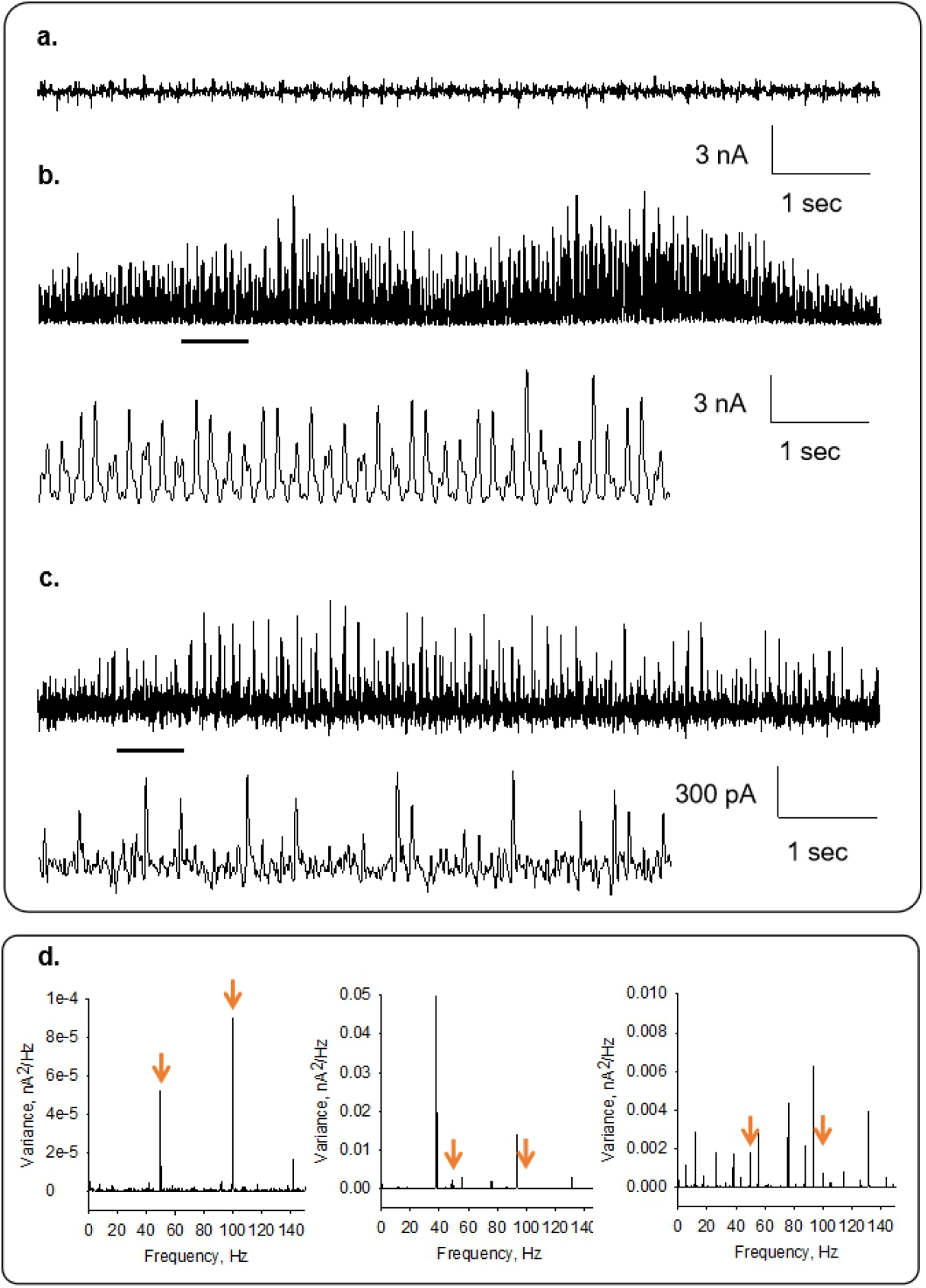
Differences in oscillatory behavior of isolated MTs and MT sheets. Representative recordings from the pipette in solution (**a**), a voltage-clamped MT sheet (**b**) and an isolated MT (**c**). Expanded tracings are also shown. (**d**) Linear-linear plots of Fourier power spectra obtained from unfiltered current responses of tracings in (**a**) (Left), (**b**) (Middle), and (**c**) (Right).

Previous studies with taxol-stabilized MTs, originally demonstrated the propagation and amplification of electrical pulses, but not the presence of electrical oscillations (8). In those studies, the MT was able to amplify both the electrical pulse injected at the stimulus as well as the collection sites, with transfer amplification ratios up to 2.35 under high ionic strength conditions (8). Although these findings demonstrated electrical amplification, oscillatory responses were never observed. In retrospect, the use of taxol to stabilize the MTs and hold them to the pipettes (8, 9) may have caused this behavior (lack thereof). Indeed, electrical oscillations of both MT sheets and bundles were readily inhibited by addition of taxol to the preparations (10, 11). Thus, herein, we explored non-taxol stabilized MTs, which instead made it more difficult to find and handle the isolated MTs in the preparation. In the present study, the electrical behavior of isolated MTs was different to that of the MT sheets present in the preparation. In particular, a broader spectrum of fundamental frequencies was always observed in isolated MTs as compared to the MT sheets. Therefore, more structured MT complexes (i.e. bundles, sheets) may render more coherent responses at given oscillatory frequencies and raise the hypothesis that combined MTs may tend to oscillate and entrain together. Different oscillatory modes have been postulated for MT structures (14, 15).

In conclusion, the present study provides to our knowledge the first experimental evidence for electrical oscillations of single brain MTs. Although the oscillating behavior has been observed in other MT structures (10, 11), the electrical behavior of isolated MTs was somewhat richer, and consistent with the presence of several fundamental frequencies that cancel out and disappear to offer more coherent behavior in brain MT sheets. MT electrical oscillations in the neuronal environment may provide a novel means for interactions between different cellular organelles and/or cytoskeletal structures as we recently observed in the context of interactions between MT sheets and actin filaments (16). Finally, the present study sheds new light into the structural/functional correlates of MT electrical oscillations (Fig. 5). Tubulin dimers, arrange in necklace-type sequences known as protofilaments, which either curl into MTs, or attach to each other laterally to form a 2D-sheets (17). Although we speculated that lateral apposition of protofilaments spontaneously extends to larger flat surfaces (10), it may have been possible for MTs to assemble and form different super structures such as sheets, bundles, or macrotubes, the possibility also exists for 2D structures from pre-formed MTs to electrostatically interact and attach laterally as well as by surface apposition (18, 19), to form different electrically-active structures instead.

**Fig 5:**
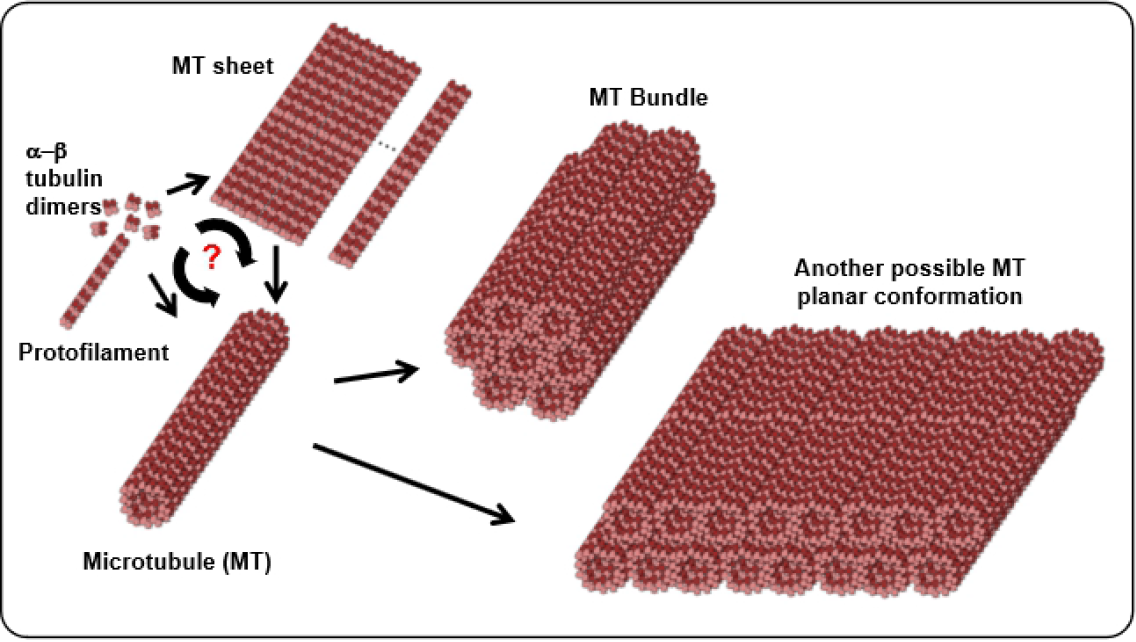
Diagram of different tubulin assembly conformations during the process of MT formation. The αβ tubulin dimers (*Left*), arrange in a necklace-type sequence known as a protofilament, which could either curl, or attach to each other to form a MT sheet, which then curves into an MT. Often times lateral apposition of protofilaments spontaneously extends to larger flat surfaces. It was originally postulated that these structures were those used in our recent patch-clamping studies (10). However, MTs may also assemble to form different super structures, including sheets, bundles, or macrotubes, such that the possibility also exists for 2D structures to occur from pre-formed MTs to attach laterally as well as by surface apposition.

## Author Contributions

BCG conducted all the experimental work and analyzed the data. MRC and HFC designed all the experiments, analyzed data and wrote the manuscript. All authors approved the final version of the manuscript.

## Acknowledgements

The authors wish to acknowledge the Ministerio de Ciencia, Tecnología e Innovación Productiva de la Nación (Argentina) for funding the studies through PICT 2016-3739.

